# Pancreatic Tumor Eradication *via* Selective PIN1 Inhibition in Cancer Associated Fibroblasts and T Lymphocytes Engagement

**DOI:** 10.1101/2022.01.31.478589

**Authors:** Jiaye Liu, Meng Li, Kewei Li, Yang Wang, Shan Li, Wenshuang Wu, Lingyao Du, Chunyang Mu, Xiaoyun Zhang, Chuan Li, Wei Peng, Junyi Shen, Yang Liu, Dujiang Yang, Kaixiang Zhang, Qingyang Ning, Xiaoying Fu, Yu Zeng, Yinyun Ni, Qiuwei Pan, Zongguang Zhou, Yi Liu, Yiguo Hu, Tianfu Wen, Zhihui Li, Yong Liu

**Author notes:** **CORRESPONDENCE**, <u>Tianfu Wen</u>, Prof. MD., PhD, Department of Liver Surgery & Liver Transplantation Center, West China Hospital, Sichuan University, room 7-1, Guoxue No.1 Road, Chengdu, China.; <u>Zhihui Li</u>, Prof. MD., PhD, Department of Thyroid and Parathyroid Surgery, West China Hospital, Sichuan University, room 5-3, Guoxue No.1 Road, Chengdu, China.; <u>Yong Liu</u>, Prof. MD., PhD, Department of Gastrointestinal Surgery, West China Hospital, Sichuan University, room 8-3, Guoxue No.1 Road, Chengdu, China.

## Abstract

Cancer associated fibroblasts (CAFs) support tumors *via* multiple mechanisms, including maintaining the immunosuppressive tumor microenvironment and limiting infiltration of immune cells. The prolyl isomerase PIN1, whose overexpression in CAFs hasn’t been fully profiled yet, plays critical roles in tumor initiation and progression. To decipher effects of selective PIN1 inhibition in CAFs on pancreatic cancer, we formulate DNA-barcoded micellular systems (DMS) encapsulating PIN1 inhibitor. DMS functionalized with CAFs-targeting antibodies (antiCAFs-DMS) can selectively inhibit PIN1 in CAFs of the tumor, leading to efficacious but temporal tumor inhibitions. We further integrate DNA aptamers (AptT), which can engage CD8^+^ T lymphocytes, to antiCAFs-DMS and thus prepare the bispecific antiCAFs-DMS-AptT system. AntiCAFs-DMS-AptT shows its potent capacity to eradicate pre-established subcutaneous and orthotopic pancreatic cancer on mice.

## INTRODUCTION

Cancer immunotherapies, including checkpoint blockade, adoptive cellular therapy, cancer vaccinology and bispecific T cell engagers, have shown potentials for cure of cancers^1–3^. Nonetheless, only a small subset of the patients within a large cohort can respond favorably to these immunotherapies^4,5^. Accumulative findings have corelated this with the immunosuppressive tumor microenvironment (TME)^6–8^, to which cancer associated fibroblasts (CAFs) greatly contribute^9–12^. CAFs are heterogeneous stromal cells and are prominent components of the TME in solid tumors and shape the immune ecosystem of TME toward a tolerant and immunosuppressive milieu via multiple mechanisms, including production of multiple cytokines and chemokines that then mediate the recruitment and functional differentiation of innate and adaptive immune cells^7,13,14^. Furthermore, CAFs can directly interact with tumor-infiltrating immune cells in negative ways, abrogating their function of tumor cell killing^15,16^.

Prolyl isomerase PIN1, which highly expresses in both cancer cells and CAFs^17^, facilitates multiple cancer-driving pathways through regulating the conformational transformation of phosphorylated Serine/Threonine-Proline motif^18,19^. Pharmacological inhibition of PIN1 has therefore been regarded as a potent anticancer strategy^17^. A few small molecule compounds that inhibit PIN1, such as all-trans retinoic acid, arsenic trioxide, juglone, AG17724, KPT-6566 and sulfopin, have been identified and used or repurposed for investigating roles of PIN1 in oncogenesis^20–24^. All these compounds, when applied *in vivo,*even though a small fraction of injected amount can finally distribute into tumor and affect tumor progression *via* inhibiting PIN1 there, it is unclear how much different cell population, like CAFs, of the tumor contributes.

To figure out exclusive effects of PIN1 in CAFs on tumor progression, one strategy can be about delivering currently available PIN1 inhibitors *via* customized drug delivery systems (DDSs) to CAFs. DDSs, including these targeting CAFs, have continuously proven their preclinical successes for cellular or even subcellular targeting drug transportation^25–29^ DDSs formulated for the delivery of PIN1 inhibitor to CAFs, however, have not been developed yet. Here, we report a DNA-barcoded micellular system to deliver PIN1 inhibitor AG17724 (designated as “DMS”) (**Scheme 1**). DMS functionalized with antibodies targeting CAFs (antiCAFs-DMS) is customized to specifically deliver AG17724 to CAFs of subcutaneous and orthotopic pancreatic ductal adenocarcinoma (PDAC), thus helping us to understand functions of PIN1 in CAFs on supporting PDAC. Furthermore, antiCAFs-DMS is mounted with immune cell-recruiting DNA aptamer (antiCAFs-DMS-AptT) to engage CD8^+^ T lymphocytes for PDAC eradication.

**Scheme 1.**
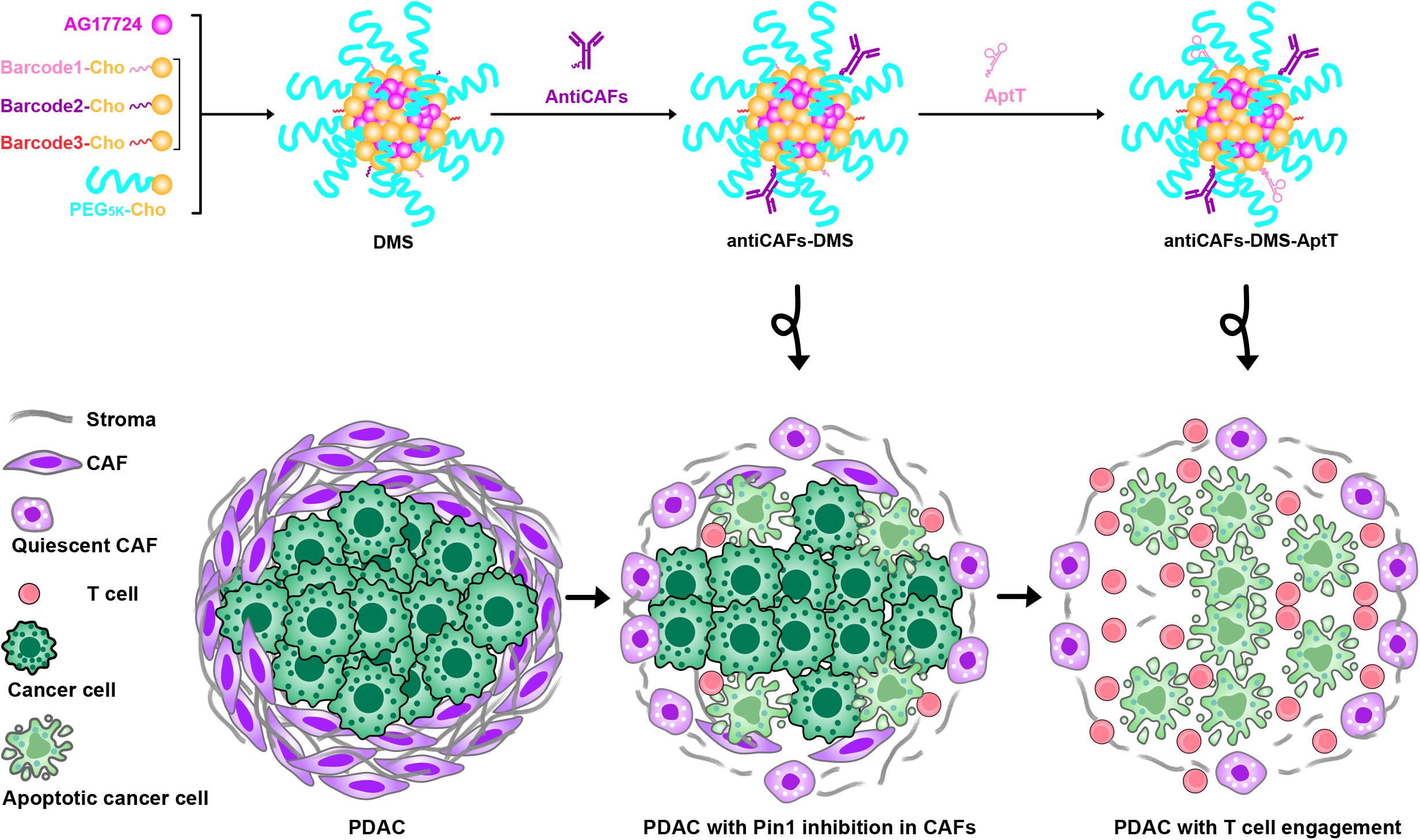
Schematic illustration of DMS, antiCAFs-DMS and antiCAFs-DMS-AptT for selective delivery of PIN inhibitor to CAFs and then directing T Lymphocytes to TME of PDAC model.

## RESULTS

### DMS packs AG17724 and displays barcodes for post functionalization

We prepared the DMS *via* self-assembly from PEG_5K_-cholesterol (PEG_5K_-Cho), DNA-barcoded cholesterol (barcodeX-Cho) and PIN1 inhibitor AG17724 (**Fig. 1A**). The hydrophobicity of AG17724 drove its encapsulation by the cholesterol core of DMS. The hydrophilic PEG_5K_ polymers surrounding DMS were supposed to stabilize the system^30^. DNA barcodes on the final DMS could facilitate post-functionalization, allowing sequential and reliable attachments of DNA-conjugated ligands. Theoretically, many different barcodes, which correspond to specific sequence of the DNA, can be used here. We used three (Table S1) to show the proof of concept. Besides, all these materials nowadays are commercially available and widely used in biomedical research, which can thus facilitate the reproductivity and accelerate the translation.

**Fig.1.**
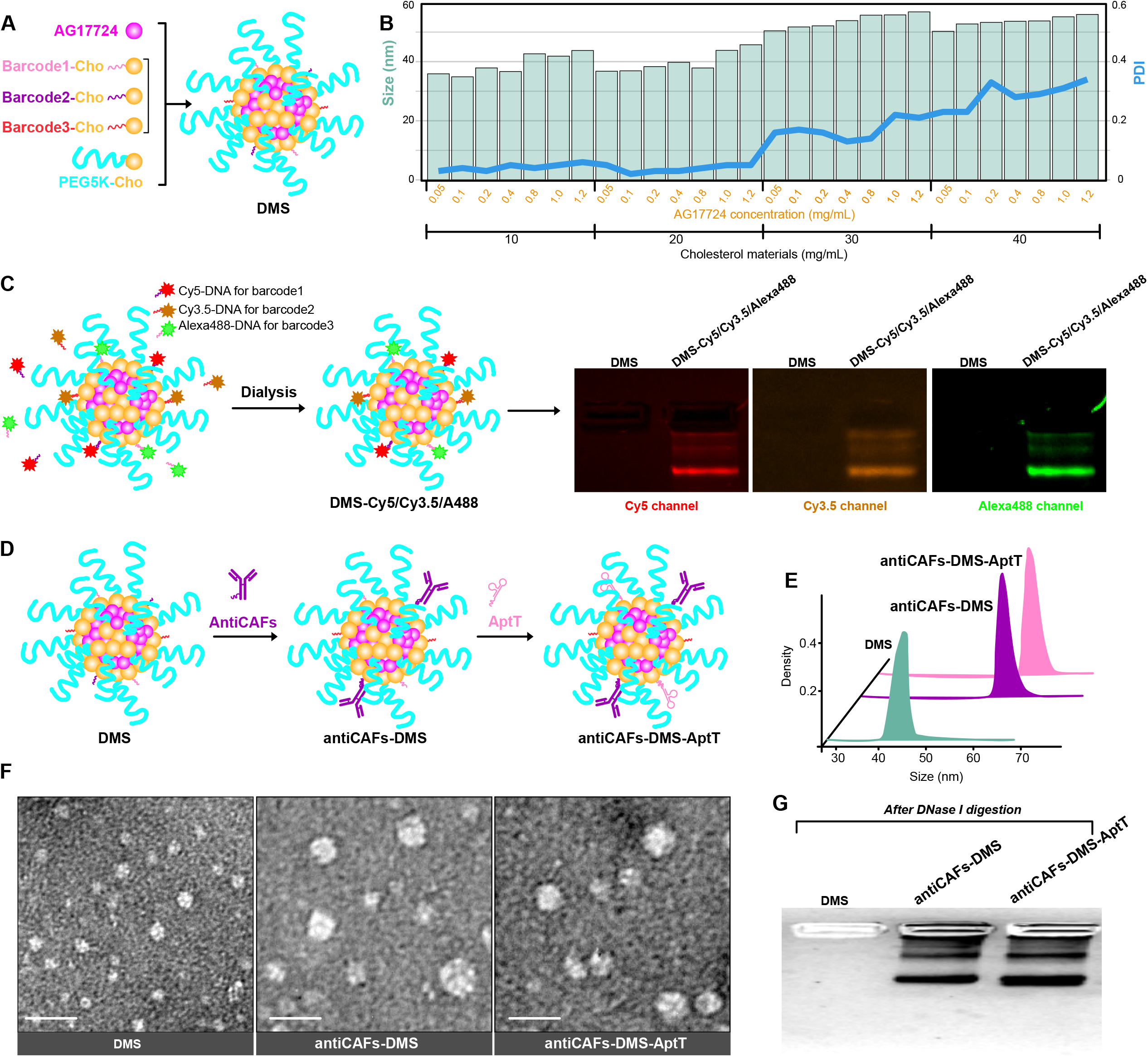
Characterization of formulations. (**A**) Schematic compositions and self-assembly of DMS. (**B**) Corresponding sizes and PDIs of DMS prepared from different concentrations of AG17724 and cholesterol materials. (**C**) Validation of DNA barcodes on DMS *via* using fluorescent probe-labeled complimentary DNA oligos. (**D**) Schematic workflow to prepare antiCAFs-DMS and antiCAFs-DMS-AptT from DMS. (**E**) Hydrodynamic size distributions of DMS, antiCAFs-DMS and antiCAFs-DMS-AptT. (**F**) TEM images of nanoparticles. Scale bars are 100 nm. (**G**) Silver staining-based antibody detections after releasing antibodies from the nanoparticles.

To figure out the optical DMS preparation, the molar ratio of barcode1-Cho/barcode2-Cho/ barcode3-Cho/PEG_5K_-Cho was fixed at 1/1/1/7, while serial concentrations of them (from 10 to 40 mg/mL) and AG17724 (from 0.05 to 1.2 mg/mL) were screened with measuring the corresponding size and polydispersity index (PDI) (**Fig. 1B**). It showed that sizes were always between 30 nm to 60 nm while PDI increased to more than 0.1 once the concentration of cholesterol materials was above 30 mg/mL. This leaded us to use “20 mg/mL cholesterol materials with 1.2 mg/mL AG17724” to prepare final DMS, which had AG17724 encapsulation efficiency and loading efficiency at 83.6 ± 1.9% and 45.8 ± 2.4%, respectively. DMS had its size at 43 ± 1.8 nm and PDI at 0.05± 0.01. DMS had a negative zeta potential at around −4.3 mV (fig. S1), which should be mainly contributed by DNA barcodes. DMS showed size stabilities with 10% serum at room temperature for one week (fig. S2). Less than 10% of loaded AG17724 was released during this one-week evaluation, further demonstrating its stability (fig. S3). This delayed release *in vitro* can reduce the risk of immediate release of AG17724 from DMS in blood stream, mitigating off-target effects. After internalizing by targeting cells, the release of AG17724 would be accelerated by the lysosomes.

To check the availability of DNA barcodes on DMS, we used complementary DNA strands attached with different fluorescent probes (Table S2), including Cy5, Cy3.5 and Alexa488, to visualize them. After removing excessive probes *via* dialysis, we ran samples on 2% agarose gel and then imaged the gels under specific channels (**Fig. 1C**). We can detect all signals from the DMS sample, indicating that all three different barcodes were well presented and accessible even though it is hard to absolutely quantify each barcode. One group of these barcodes were used as coordinates to hybridize DNA-conjugated CAFs-targeting antibodies (antiCAFs) (fig. S4) to get antiCAFs-DMS (**Fig. 1D**). We used the antibody recognizing the membrane biomarker, fibroblast activation protein alpha (FAP-α), on CAFs as the antiCAFs^31^. Furthermore, aiming to direct immune cells against tumor, we hybridized DNA aptamers (Table S3), which were screened to bind CD8^+^ T lymphocytes with high specificity and selectivity^32^, with another group of barcodes on antiCAFs-DMS to get the bispecific antiCAFs-DMS-AptT. Hydrodynamic sizes of both antiCAFs-DMS and antiCAFs-DMS-AptT were similarly around 63 nm (**Fig. 1E**), which were around 20 nm bigger than DMS. Transmission electron microscopy (TEM) imaging of negatively stained samples further validated size differences among these three preparations (**Fig. 1F**). We verified the existence of antibody on both antiCAFs-DMS and antiCAFs-DMS-AptT by gel silver staining after releasing antibody via DNase I digestion (**Fig. 1G**).

### Antibody-functionalized DMS selectively inhibits PIN1 in CAFs

We cultured CAFs *via* inducing the differentiation of normal fibroblasts (NIH-3T3 cells) with transforming growth factor beta (TGF-β)^33^, and we validated its reliability *via* showing drastically increased expressions of both FAP-α and alpha-smooth muscle actin (α-SMA) (**Fig. 2A**). Flow cytometry-based cellular uptake analysis showed the high selectivity of both antiCAFs-DMS and antiCAFs-DMS-AptT on CAFs rather than pancreatic cancer cells (Pan02), displaying around 70-fold differences (**Fig. 2B**). Further qualitative (**Fig. 2C**) and quantitative (**Fig. 2D**) analysis on the uptake of antiCAFs-DMS among Pan02 cells, NIH-3T3 cells and CAFs demonstrated the similar differences.

**Fig.2.**
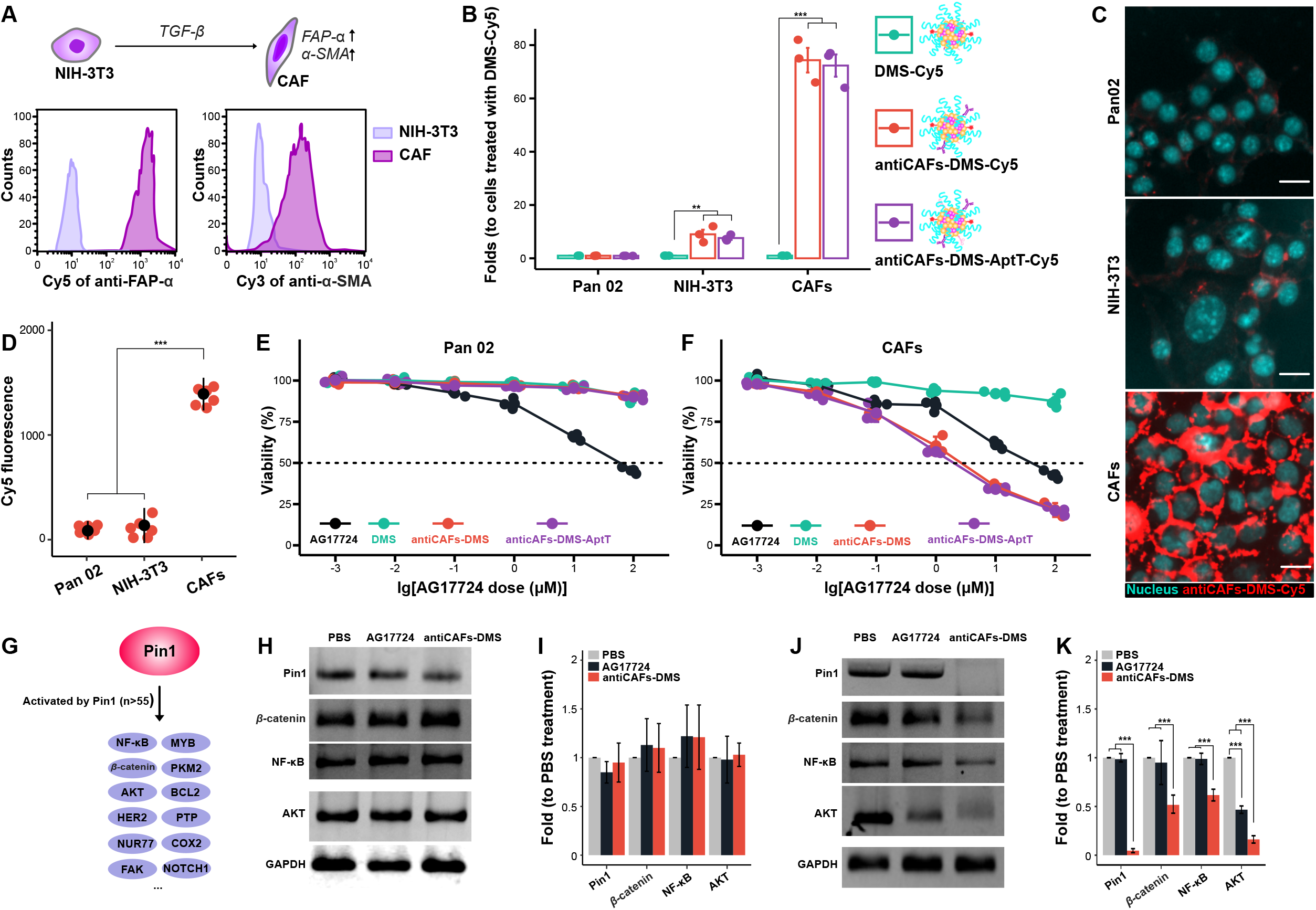
Specific PIN1 inhibition in CAFs achieved by antibody modified DMS *in vitro*. (**A**) Establishment and validation of CAFs *via* high expressions of both FAP-α and α-SMA. (**B**) Cellular uptake of Cy5-labeled formulations on Pan02 cells, NIH-3T3 cells and CAFs, measured by flow cytometry. (**C**) Cellular uptake of Cy5-labeled antiCAFs-DMS on cells, imaged by fluorescent microscopy. Scale bars are 20 μm. (**D**) Cellular uptake of Cy5-labeled antiCAFs-DMS on cells, quantified by flow cytometry. (**E**) Cytotoxicity on Pan02 cells and CAFs (**F**). (**G**) PIN1 regulates multiple cancer-related pathways. (**H**) Western blotting detection of PIN1 and relative proteins from Pan02 cells treated with PBS, AG17724 or antiCAFs-DMS. (**I**) Quantitative analysis of the western blotting results from Pan02 cells. (**J**) Western blotting detection of PIN1 and relative proteins from CAFs treated with PBS, AG17724 or antiCAFs-DMS. (**K**) Quantitative analysis of the western blotting results from CAFs.

We then studied the cytotoxicity of all formulations on Pan02 cells (**Fig. 2E**) and CAFs (**Fig. 2F**). Owe to its poor cell permeability^34–36^, compound AG17724 itself showed high IC_50_ at around 50 μM for both cells. IC_50_ of DMS exceeded our experimental range for both cells, which was due to its low cell uptake efficiency. Intriguingly, antiCAFs-DMS and antiCAFs-DMS-AptT showed their IC_50_ on CAFs at around 1.2 μM, whereas their cytotoxicity on Pan02 cells were much lower. This indicated that attached FAP-α antibodies can firstly help antiCAFs-DMS and antiCAFs-DMS-AptT targeting CAFs, they then also can facilitate the internalization of the whole systems by cells, achieving a specific and efficient delivery of PIN1 inhibitor into CAFs.

To see if PIN1 could be inhibited, cells were treated with 0.5 μM of AG17724 for 4 hours and relative proteins were analyzed. As a critical modifier of multiple cancer-related pathways, PIN1 has been revealed to activate more than 55 proteins, including β-catenin, NF-ºB and AKT (**Fig. 2G**). We herein checked them, and we observed that, for pancreatic cancer cells, PIN1 and these three relative proteins were at the similar levels no matter they were treated with phosphate buffered saline (PBS), AG17724 or antiCAFs-DMS (**Fig. 2H, 2I**). For CAFs, antiCAFs-DMS significantly inhibited PIN1, also efficiently inhibited β-catenin, NF-ºB and AKT to different extents. AG17724 itself showed the capacity to inhibit AKT, however, it didn’t block PIN1, β-catenin and NF-ºB (**Fig. 2J, 2K**). These results showed similar patterns to the cellular uptake part and cytotoxicity part aforementioned, further indicating that antiCAFs-DMS was potent for selective PIN1 inhibition in CAFs.

### PIN1 inhibitions in CAFs reduce invasion and growth of pancreatic cancer spheroids

CAFs have been revealed to enhance growths of different tumors^11^. For pancreatic cancer, especially, the molecular crosstalk between cancer cells and CAFs facilitates tumor invasion and growth^17,37^. To determine whether selective PIN1 inhibition in CAFs affects their ability to act on pancreatic cancer cells, we conducted the indirect co-culture of pre-formed pancreatic cancer spheroids with CAFs (**Fig. 3A**). Unlike spheroids cultured under the same condition but without adding CAFs, spheroids with indirect CAFs coculture showed invasive cells at their surfaces, confirming that CAFs can remotely act on cancer cells, which could be *via* humoral factors (**Fig. 3B**). AG17724 (0.5 μM) failed to inhibit invading signs around spheroids in matrigel. In contrast, the antiCAFs-DMS treatment stopped the invasion of spheroids, displaying shrinking margins. The dynamic volume records of these spheroids further showed the inhibitive effect of antiCAFs-DMS on their growth (**Fig. 3C**). These results together firstly highlighted that PIN1 is important for CAFs to promote growth and invasion of pancreatic cancer spheroids. Secondly, we showed that PIN1 inhibitor AG17724 delivery, *via* antiCAFs-DMS, into CAFs might be an alternative and potent way for pancreatic cancer treatment.

**Fig.3.**
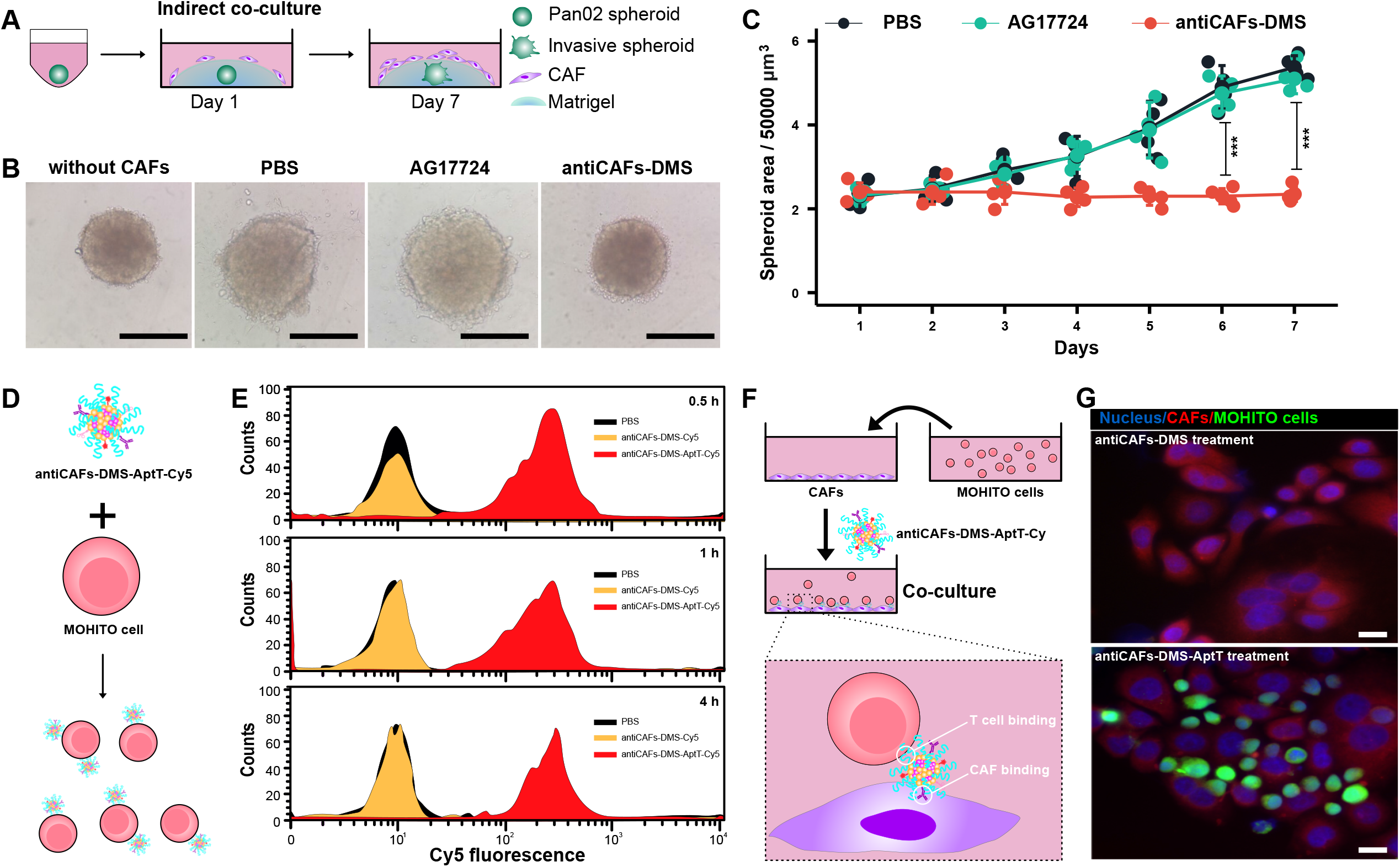
Cancer cells and T cell engagement *via* bispecific antiCAFs-DMS-AptT in 3D and 2D models. (**A**) Schematic workflow of Pan02 spheroids and CAFs coculture. (**B**) Representative morphologic images of Pan02 spheroids at the end of the indirect coculture. Scale bars are 100 μm. (**C**) Dynamic growth curves of Pan02 spheroids during the indirect coculture. (**D**) Schematic workflow of the incubation of MOHITO cells with Cy5 labeled antiCAFs-DMS-AptT. (**E**) Quantitative analysis of the binding of Cy5-labeled antiCAFs-DMS or antiCAFs-DMS-AptT to MOHITO cells *via* flow cytometry. (**F**) Schematic illustration of the direct coculture of CAFs and MOHITO cells. (**G**) Representative microscopic images of the coculture experiment. Scale bars are 10 μm.

### Aptamer modification on antiCAFs-DMS bridges T Lymphocytes to CAFs

Solid tumors, PDAC especially, often show their resistances to immunotherapy^38^. Apart from their own immunosuppressive TME, in which cancer cells debilitate the antitumor immunity of resident immune cells *via* secretion or presentation of inhibiting molecules and collaboration with CAFs, dense architectures, largely contributed by CAFs, raise physical obstacles for active peripheral immune cells to infiltrate.^39–41^ With interest of bridging active peripheral T lymphocytes to PDAC, we prepared antiCAFs-DMS-AptT as mentioned above. After incubating murine CD8^+^T cells with Cy5-labbled systems (**Fig. 3D**), cytometry analysis showed that cells treated with antiCAFs-DMS-AptT had much higher fluorescent intensity than cells treated with antiCAFs-DMS, confirming that this aptamer decoration can efficiently lead the system to stick to T cells (**Fig. 3E**). There was no big difference, in term of Cy5 signal on T cells, among time points of 0.5, 1 and 4 hours, indicating that both binding and saturation were fast.

Since antiCAFs-DMS-AptT contains antibodies for targeting CAFs but also aptamers for the binding of CD8^+^ T cells, it ideally could function as bispecific modules, which have already shown promising potentials in immunotherapy^42–44^, to bridge T cells to CAFs. To validate this, we conducted the coculture of CAFs and T cells *in vitro* (**Fig. 3F**). We observed that T cells barely located on CAFs under antiCAFs-DMS treatment. AntiCAFs-DMS-AptT treatment, nevertheless, showed its effect to bridge T cells to CAFs (**Fig. 3G**). These results indicated that, apart from the delivery of PIN1 to CAFs, antiCAFs-DMS-AptT potentially can also direct T lymphocytes to CAFs, resulting in more immune cells in TME. Being different from bispecific antibodies, aptamers or nanoparticles, which usually still can’t overcome physical obstacles of PDAC, antiCAFs-DMS-AptT can render solid tumor accessible for T lymphocytes.

### Biodistribution and effects from PIN1 inhibition in CAFs and T cells engagement

To assess if our delivery systems could successfully distribute into tumors *in vivo*, we labeled them with Alexa750 and then intravenously administrated them to mice bearing subcutaneous PDAC. After 4 hours, we can see that livers were the main bio-distributing organs of all systems (**Fig. 4A**). Very low Alexa750 signals were detected on hearts, lungs, spleens and kidneys. For tumors, it showed that both antiCAFs-DMS and antiCAFs-DMS-AptT can accumulate to tumors much more efficiently than DMS. To make it more precise and quantitative, we prepared homogenates from these collected organs or tumors, and then imaged them and performed quantitative analysis (**Fig. 4B, 4C**). It showed same trends as organ imaging, quantitively presenting that, compared with DMS, antibody decorations improved its tumor accumulation by around two folds.

**Fig.4.**
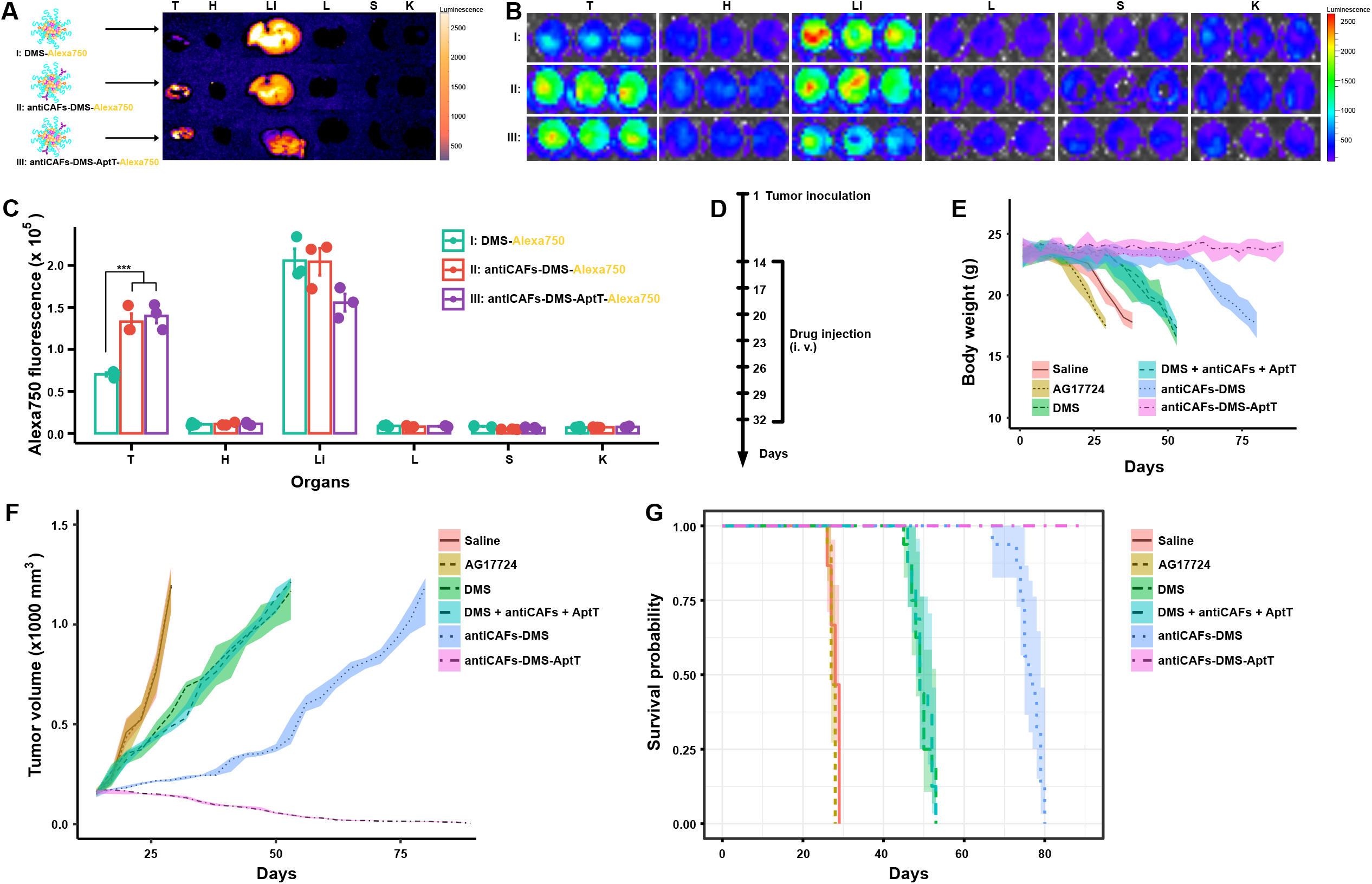
Biodistribution and therapeutic effects. (**A**) Representative *ex vivo* imaging of tumors and organs (T, tumor; H, heart; Li, liver; L, lung; S, spleen; K, kidney) from mice intravenously injected with Alexa750 labeled DMS, antiCAFs-DMS and antiCAFs-DMS-AptT. (**B**) Representative imaging of homogenates prepared from tumors or organs (T, tumor; H, heart; Li, liver; L, lung; S, spleen; K, kidney) from mice intravenously injected with Alexa750 labeled DMS, antiCAFs-DMS and antiCAFs-DMS-AptT. (**C**) Quantitative Alexa750 fluorescent intensity analysis of homogenates prepared from tumors or organs (T, tumor; H, heart; Li, liver; L, lung; S, spleen; K, kidney). (**D**) treatment schedule. (**E**) Body weight records of tumor-bearing mice receiving different treatments. (**F**) Tumor volume growth curves. (**G**) Kaplan–Meier tumor-inoculated mouse survival curves.

We then evaluated their antitumor effects *in vivo* by following our treatment schedule (**Fig. 4D**). Body weight changes were recorded. Compared to control mice (saline), AG17724 injection (10 mg/kg) accelerated weight loss, proving that the compound itself caused toxicity to mice. All treatments with nano-formulated AG17724 slowed down the loss of body weight to different extents. AntiCAFs-DMS-AptT treatment showed its capacity to stabilize body weight during the whole experimental period (**Fig. 4E**). On tumor growth, we didn’t see any effect of AG17724 itself. Both DMS and the mixture of DMS, antibody and aptamers (DMS + antiCAFs + AptT) administrations slightly delayed the tumor growths. AntiCAFs-DMS stabilized tumors for around 40 days, but it failed to inhibit their growths afterwards. AntiCAFs-DMS-AptT showed its promising anti-tumor capacity, almost eradicating the established tumors (**Fig. 4F**). Accordingly, survival time of mice also ranked antiCAFs-DMS-AptT as the best anti-tumor formulation among our treatments (**Fig. 4G**).

### CAFs and T cells quantification in tumor tissues

To check if our CAFs-targeting PIN1 inhibitor delivery systems worked, as expected, to block PIN1 in tumors during the treatment, we took tumors at the 25^th^ day for fluorescence-activated cell sorting analysis. CAFs counting showed us that percentages of CAFs inside tumors were reduced from 36.3% to 20.3%, 5.6% and 2.5% by DMS, antiCAFs-DMS and antiCAFs-DMS-AptT, respectively (**Fig. 5A, 5B**). This indicated that DMS functionalized with CAFs-targeting antibody can improve its specificity, resulting in more CAFs depletion.

**Fig.5.**
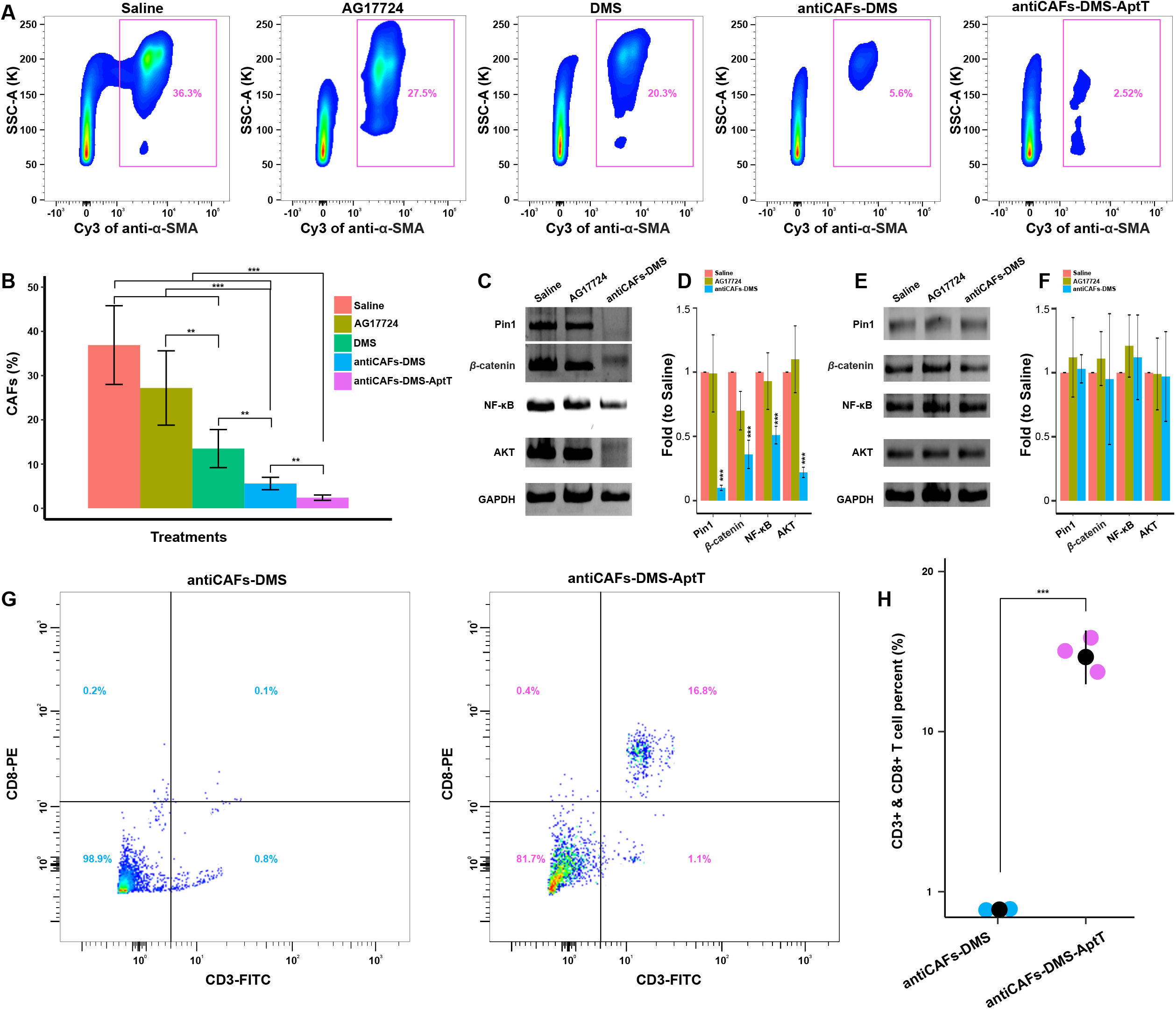
Tumor tissue analysis on CAFs, PIN1 and T lymphocytes. (**A**) and (**B**) Quantitative analysis, via cell sorting and counting, of CAFs in tumors from mice receiving different treatments. (**C**) Western blotting detection of PIN1 and relative proteins in sorted CAFs from tumors collected from mice treated with saline, AG17724 or antiCAFs-DMS. (**D**) Quantitative analysis of the western blotting results of sorted CAFs. (**E**) Western blotting detection of PIN1 and relative proteins in non-CAFs cells from tumors collected from mice treated with saline, AG17724 or antiCAFs-DMS. (**F**) Quantitative analysis of the western blotting results of non-CAFs cells. (**G**) and (**H**) Quantitative analysis, via cell sorting and counting, of CD8^+^ T lymphocytes in tumors from mice treated with antiCAFs -DMS or antiCAFs-DMS-AptT.

PIN1 and related proteins in these sorted cells were analyzed *via* western blots. For CAFs in tumors, PIN1 was significantly inhibited by antiCAFs-DMS treatment whereas AG17724 didn’t display its effect. Besides, β-catenin, NF-_κ_B and AKT, whose activations are partially controlled by PIN1, were also inhibited by antiCAFs-DMS (**Fig. 5C, 5D**). For sorted pancreatic cancer cells from the tumors, we didn’t see amount changes of these proteins (**Fig. 5E, 5F**). This directly and further confirmed the delivery specificity of antiCAFs-DMS towards CAFs in tumors. Apart from PIN1 inhibition in CAFs, antiCAFs-DMS-AptT treatment, which almost eradicated the established tumors, was also supposed to function as bispecific CD8^+^ T cells engager. We proved this via comparing CD8^+^ T cells populations inside tumors treated with antiCAFs-DMS or antiCAFs-DMS-AptT. We detected around 16% more intratumor CD8^+^ T cells (16.8%) under antiCAFs-DMS-AptT treatement than antiCAFs-DMS treatment (0.1%) (**Fig. 5G, 5H**). This confirmed our expectation, also told us that the combination of PIN1 inhibition in CAFs and immune cell engaging could be a potent way to render PDAC eradicable.

### Response of orthotopic murine pancreatic cancer to antiCAFs-DMS-AptT

We further established orthotopic PDAC to assess if antiCAFs-DMS-AptT treatments still eradicate cancer cells in pancreas. Luciferase-expressing Pan02 cells (Pan02-Luc) we used here helped us, *via* direct bioluminescent imaging, monitoring the cancer developments. Mice received their first treatment at the 14^th^ day post cancer cell implant (**Fig. 6A**). Dose administration of each formulation corresponded to 10 mg/kg AG17724. It showed that, after four rounds of treatments, mice of antiCAFs-DMS-AptT group had almost no bioluminescent signals from cancer cells (**Fig. 6B**), indicating the potent anti-PDAC efficacy of antiCAFs-DMS-AptT on orthotopic pancreatic cancer model. Although DMS or antiCAFs-DMS also showed their capacity to slightly slow down the growth rate of cancer cells in pancreas, mice treated with them experienced increasing tumor burdens during the period of experiment (**Fig. 6C**). Correspondingly, antiCAFs-DMA-AptT significantly prolonged the survival time of mice, and the survival rate was 100% during our 80-day investigation (**Fig. 6D**).

**Fig.6.**
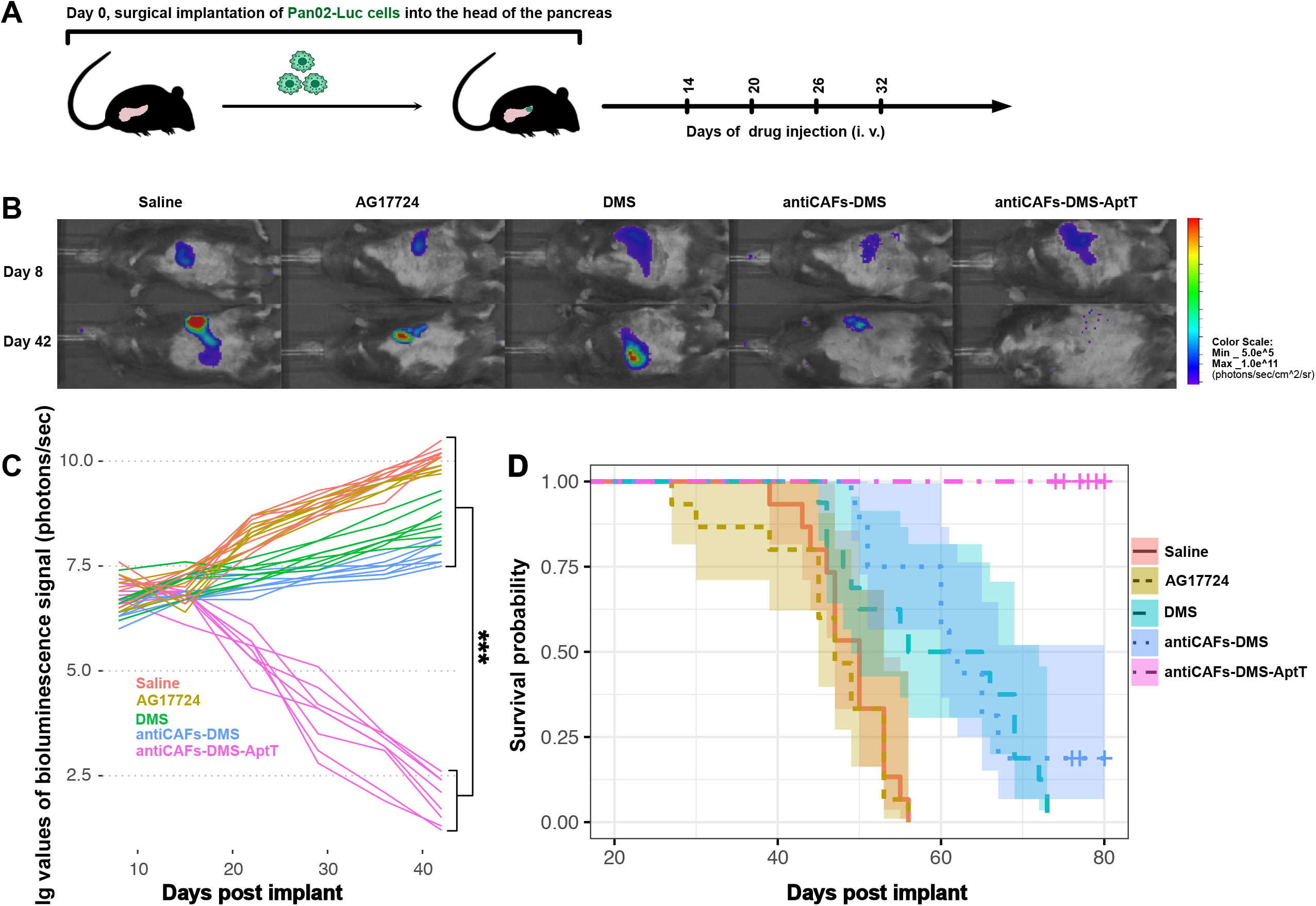
Treatment effects of antiCAFs-DMS-AptT on orthotopic pancreatic cancer model. (A) Treatment schedule of mice bearing orthotopic pancreatic cancer. (B) Representative bioluminescent images of mice treated with different formulations at the day 8 and day 42 post cancer establishment. (C) Tumor development of each mouse, quantified by bioluminescence signal. (D) Kaplan-Meier survival curves of mice.

## DISCUSSION

Due to the critical roles of PIN1 in tumor initiation and progression, small molecules for inhibiting PIN1 have continuously been screened or developed. However, these inhibitors can lose pharmacological activity they had *in vitro* once administrated and diluted *in vivo*. Besides, a few available drugs have been repurposed for PIN1 inhibition in tumors, which can generally inhibit PIN1 but can’t restrict the inhibition within specific cell population of the tumor^17,20–24^. The DNA-barcoded micellular systems we are developing in this work prove the feasibility to regulate PIN1 in cell population of interest. This will then give insights into cell-level antitumor targeting therapy rather than at the whole tumor tissue level.

Densely structured solid tumors are usually resistant to current therapies, which are highly contributed by CAFs^14,41^. Recent literature showed that overexpressed PIN1 displays both in cancer cells and CAFs and aggravates PDAC^17^. How much PIN1 in CAFs contributes to tumor progression, however, is unknown since the lack of drugs exclusively blocking PIN1 in CAFs. We find that PIN1 inhibitor AG17724 can be targeted into CAFs, both *in vitro* and *in vivo*, via delivering it by antiCAFs-DMS. Functionally, antiCAFs-DMS selectively and drastically inhibited PIN1 in CAFs, resulting in higher cytotoxicity on CAFs than pancreatic cancer cells. AG17724 itself didn’t show any antitumor therapeutic effect *in vivo*, which might be related to the poor cell permeability of the compound^34–36^. Intriguingly, we find that effective PIN1 inhibition in CAFs, via antiCAFs-DMS, can slow down the growth of both subcutaneous and orthotopic pancreatic PDAC. However, it is not a potent way to stop tumors since they grew fast once we halted drug injections.

Immunotherapies could be combined with chemotherapy to get better antitumor efficacy^45^, which inspired us to prepare our bispecific antiCAFs-DMS-AptT. Artificial bispecific modules, including bispecific antibody, bispecific aptamers and bispecific nanoparticles, are emerging as potent antitumor drugs via directing immune cells of host against cancer cells^42–44^. The highly desmoplastic TME of certain types of solid tumors, however, can limit their ability to bring immune cells into tumor tissue^6^. We find that antiCAFs-DMS-AptT can eradicate the established tumors. We speculate that PIN1 inhibition in CAFs by antiCAFs-DMS-AptT firstly can disrupt the desmoplastic and immunosuppressive TME, leading to an accessible TME for immune cell infiltration. Secondly or simultaneously, T lymphocytes engaged by antiCAFs-DMS-AptT can, more easily, distribute into the tumor, initiating tumor eradication.

In summary, we demonstrated antibody-functionalized DNA-barcoded micellular systems by which selective PIN1 inhibition in CAFs can be achieved. This renders pancreatic tumors eradicable by cytotoxic T lymphocytes engagement. DNA barcodes on these micellular systems offer an easy-to-change way to customize the targeting demand, opening the way for PIN1 inhibition in certain cell populations of the tumor and thus uncovering mechanistic details on how PIN1 supports tumor initiation and progression.

## METHOD

### Ethical regulation

All animal experimental procedures were approved by the Animal Experimentation Ethics Committee of Sichuan University.

### Reagents, cells and mice

AG17724, absolute ethanol, chloroform and dimethyl sulfoxide were purchased from Sigma-Aldrich (CA, USA). Cholesterol-DNA conjugates, florescent probe modified DNA oligonucleotides, DBCO-modified DNA oligonucleotides and DNA aptamers were synthesized by Integrated DNA Technologies (CA, USA). PEG_5K_-Cho was purchased from Nanosoft Polymers (Winston-Salem, USA). Dulbecco’s Modified Eagle Medium (DMEM, high glucose), fetal bovine serum (FBS), penicillin-streptomycin, Trypsin-EDTA (0.25%) and phosphate-buffered saline (PBS) were purchased from Thermo Fisher SCIENTIFIC (MA, USA). A mouse anti-FAP-*α* antibody (MAB9727-100) was purchased from R&D SYSTEMS (Minneapolis, USA), a mouse anti-*α*-SMA antibody (ab5694), a mouse anti-PIN1 antibody (EPR18546-317), a mouse anti*β*-catenin antibody (IGX4794R-3), a mouse anti-NF-ºBantibody (p65(NFkappaB)), a mouse anti-AKT antibody (phospho T308), a mouse anti-GAPDH antibody (6C5), a Cy3 conjugation kit for antibody labeling (ab188287) and a Cy5 conjugation kit for antibody labeling (ab188288) were purchased from abcam (Cambridge, UK). An HRP-conjugated goat anti-mouse IgG (H+L) cross-adsorbed secondary antibody (G-21040), a FITC-labeled mouse anti-CD3 antibody (17A2) and a PE-labeled mouse anti-CD8 antibody (12-0081-82) were purchased from Thermo Fisher SCIENTIFIC (MA, USA). Mouse TGF-*β* was purchased from Peprotech (New Jersey, USA). Pan02 cell line and NIH-3T3 cell line were purchased from ATCC (USA). Pan02-Luc cell line was purchased from Labcorp (USA). Immortalized mouse CD4^+^ CD8^+^ T cell line (MOHITO) were purchased from abm (USA). Female C57BL/6□mice were purchased from Chengdu Dashuo Biological Institute (Chengdu, China).

### Preparation and characterization of nano-formulations

We used the thin-film hydration method as described in our previous study to prepare DMS. In 100-mL round-bottomed flasks, series amounts (as indicated in Fig. 1B) of barcode1-cho, barcode2-cho, barcode3-cho, PEG_5K_-Cho (the molar ratio of barcode 1-cho/barcode2-cho/ barcode3-cho/PEG_5K_-Cho was fixed at 1/1/1/7) and AG17724 were dissolved in organic solvent containing 10 mL of absolute ethanol and 1mL for 1 hour under 37 °C. To obtain thin films, under vacuum, the solvent was removed using rotary evaporation. The flasks were further dried in vacuum drier overnight. The thin films were hydrated in PBS for 0.5 hour in a 37°C water bath. Collected solutions were sonicated by a probe sonicator at 80 W for 75 seconds. The last step was to filter samples through sterile polyethersulfone membranes with pore size at 0.22 μm (Sigma-Aldrich, USA).

To prepare antiCAFs-DMS, DMS was incubated with DNA-antibody conjugate, whose molar amount was 10-fold excessive than barocode1 on DMS, in PBS at 37 °C overnight. The sample was centrifugated at 2K g for 5 minutes, excessive DNA-antibody conjugates were in the supernatant and was removed.

To prepare antiCAFs-DMS-AptT, DMS was incubated with DNA-antibody conjugate (10-fold excessive than barocode1 on DMS) and DNA apatamers (10-fold excessive than barocode2 on DMS) in PBS at 37 °C overnight. The sample was centrifugated at 2K g for 5 minutes, excessive DNA-antibody conjugates and DNA aptamers were in the supernatant and were then removed.

Particles sizes, PDI and zeta potentials were measured via Malvern Zetasizer Nano ZS90 instrument (Malvern Instruments Ltd., U. K.).

### Drug release assay

*In vitro* AG17724 release study was performed using a dialysis method at 37°C for 1 □ week. PBS containing 0.1% (v/v) Tween 80 was used as the release buffer. Nano-formulations were placed into dialysis tubes (MWCO = 8000 Da) and tightly sealed. Then the dialysis tubes were placed into 40 mL of release buffer and were incubated under 37 °C with gentle oscillating at 50 rpm. At specific time points, 1 mL of samples were taken, then centrifuged for 0.5 hour. AG17724 released was quantified in the supernatant by HPLC. 100 μL of release medium was taken out and replaced with equal volume of fresh release buffer. Then the samples were diluted with dimethyl sulfoxide and the concentrations of AG17724 were determined at the wavelength *λ* = 225 nm by HPLC.

### DNA-antibody conjugation

We used SiteClick^™^ Antibody Azido Modification Kit (Invitrogen, USA) to conjugate DNA oligos to the Fc region of IgG antibody. Mouse anti-FAP-*α* antibody was concentrated and buffer exchanged to 200 μg in 50 μL antibody preparation buffer (Component A). We then modified the carbohydrate domain of the antibody via adding 10 μL of β□galactosidase (Component D) and 6-hour incubation. Azide-attached antibody was produced via an overnight reaction (30 °C) with UDP-GalNAz (Component E) and GalT enzyme (Component H). At the last step, DBCO-modified DNA oligos were added to the azide-attached antibody for 4-hour copper-free click chemistry action. During each step, the product of interest was purified via 50K Amicon Ultra-0.5 mL Centrifugal Filters.

### Cell culture

Pan 02 cells and NIH-3T3 cells were cultured in DMEM (high glucose) medium containing 10% FBS, 50 units/mL of penicillin and 50 μg/mL of streptomycin at 37 °C in a 5% CO_2_ humidified environment incubator (Thermo Fisher Scientific, USA).

To get CAFs, NIH-3T3 cells were incubated with 10 ng/mL of TGF-β for 24 hours. To validate CAFs, cells suspensions were collected and incubated with Cy5-labeled anti-FAP-α antibody or Cy3-labeled anti-α-SMA antibody under room temperature for 20 minutes. After washing with cold PBS for 3 times, 50K cells were analyzed by flow cytometer (Cytomics^™^ FC 500, Beckman Coulter, Miami, FL, USA) to quantify the fluorescent intensity.

MOHITO cells were cultured in 6-well plate at 37 °C in a 5% CO_2_ humidified environment incubator (Thermo Fisher Scientific, USA). The medium for this cell line was Prigrow II medium (abm, USA) containing 20% FBS, 10 ng/mL mouse IL-7, 50 units/mL of penicillin and 50 μg/mL of streptomycin.

### Cellular uptake assays

For quantitative analysis, cells were seeded into six-well plates at the density of 100K cells per well and cultured for 24 hours. Cy5-labeled DMS, antiCAFs-DMS or antiCAFs-DMS-AptT were added to cells, at the final Cy5 concentration of 2 mg/mL, for 4-hour incubation. Cells were washed with cold PBS twice, trypsinized and resuspended in 0.5 mL of PBS. The Cy5 intensity of cells was measured by a flow cytometer (Cytomics^™^ FC 500, Beckman Coulter, Miami, FL, USA). 50K cells were recorded for each sample.

For microscopic imaging, cells, at the density of 10K cells per well, were seeded into 8-well chamber (Millicell^®^ EZ SLIDES, Merck Millipore) 24 hours prior to adding our nano-formulations. After 4-hour incubation, cells in wells were washed with cold PBS for three times and followed by fixing in 4% (v/v) paraformaldehyde. Nucleus were stained with Fluoroshield Mounting Medium with DAPI (Abcam). Fluorescence imaging was performed on Axio Imager.M2 (Zeiss, Germany).

### Cell viability assay

We performed ATP-based luminescent cell viability assay using CellTiter-Glo Luminescent Cell Viability Assays (Promega). Cells were seeded into 96-well opaque white polystyrene microplate (Corning), at the density of 20K cells per well, and cultured for 24 hours. Cells were incubated with various concentrations of AG17724, DMS, antiCAFs-DMS or antiCAFs-DMS-AptT. After 48 hours, plates were equilibrated at room temperature for 0.5 hour. Cells were lysed in 100□μl, of CellTiter-Glo reagent (Promega) for 2 minutes. After incubating under room temperature for 10 minutes, the luminescence was measured on a multimode microplate reader (Thermo Fisher Scientific Varioskan Flash, USA).

### Western blotting

Cells of interest were harvested and lysed by cell lysis buffer (Beyotime, China). Electrophoresis of samples was run on home-made 10% SDS-PAGE gels. Then samples were transferred from the gel to polyvinylidene fluoride (PVDF) films, which were next incubated with anti-primary antibodies (1:1K) for 24 overnight. PVDF films were incubated with HRP-labeled goat anti-rabbit secondary antibodies (1:5K) and washed. At the last, secondary antibodies were detected by Immobilon Western HRP Substrate (Millipore, Billerica, USA) on ChemiScope 6000 Touch System (Shanghai, China).

### Indirect coculture of pancreatic cancer spheroids and CAFs

To culture pancreatic cancer spheroids, at 37 °C in a 5% CO2 humidified environment incubator (Thermo Fisher Scientific, USA), 3K Pan 02 cells (per well) were seeded into Ultra Low Attachment 96-well plate with round bottom (Corning) for 7-day culture in DMEM (high glucose) with 20% FBS, 50 units/mL of penicillin and 50 μg/mL of streptomycin.

For indirect co-culture experiment, 10 organoids were transferred into 50-mL GFR Matrigel (356231, Corning) in DMEM (high glucose) with 20% FBS, 50 units/mL of penicillin and 50 μg/mL of streptomycin. 50K CAFs, which were pre-incubated with 0.5 μM of AG17724 or antiCAFs-DMS, were then seeded on the top of Matrigel for 7 days, followed by recording organoid growth using Cyntellect Celigo (Cyntellect) and analyzing the organoid size using Cyntellect Celigo software (version 1.3, Cyntellect).

### CD8^+^ T cell binding assay

200K of MOHITO cells per well in six-well plate were cultured for 24 hours. Cy5-labeled antiCAFs-DMS or antiCAFs-DMS-AptT were added to cells, at the final Cy5 concentration of 2 mg/mL, for 0.5-hour, 1-hour or 4-hour incubation. After washing with cold PBS twice, cells were resuspended in 0.5 mL of PBS. The Cy5 intensity of cells was measured by a flow cytometer (Cytomics™ FC 500, Beckman Coulter, Miami, FL, USA).

### Confocal microscopy study of cell–cell interaction

10K of CAFs per well were seeded into 8-well chamber (Millicell^®^ EZ SLIDES, Merck Millipore) for overnight culture and then stained by CellTracker^™^ Red CMTPX Dye (Invitrogen, USA) and Hoechst (Invitrogen, USA) for 10 minutes at 37 °C. In parallel, 50K of MOHITO cells per well in six-well plate were cultured for overnight culture and then stained by CellTracker^™^ Green CMFDA Dye (Invitrogen, USA) and Hoechst (Invitrogen, USA) for 10 minutes at 37 °C. Washing CAFs and MOHITO cells with corresponding cell culture medium (4 °C) to remove the unstained solution. MOHITO cells were then added to CAFs for coculture, and they were treated with antiCAFs-DMS or antiCAFs-DMS-AptT (control the concentration of AG17724 at 0.1 μM) at 4 °C for 2 hours. Then, cells were fixed with paraformaldehyde at a final concentration of 1% (w/v), and the images of cell–cell complexes were observed by confocal microscopy (TCS SP5 AOBS confocal microscopy system, Leica, Germany).

### Subcutaneous PDAC model, biodistribution and antitumor assay

We established the subcutaneous PDAC model by subcutaneous inoculation (at the right back of C57BL/6 mice) of 100-μL mixed cell suspension containing 1000K Pan 02 cells and 500K NIH-3T3 cells. After 2 weeks, tumors grew to the volume of around 180 mm^3^.

For biodistribution imaging, nine subcutaneous PDAC-bearing mice were randomly divided into three groups. Alexa750-labeled DMS, antiCAFs-DMS and antiCAFs-DMS-AptT were intravenously injected at 200 mg/kg of Alexa750. After 4 hours, mice were euthanized and tumors and organs were imaged using the IVIS Spectrum system (Caliper Life Sciences, Hopkinton, MA, USA). Then, tumors and organs were homogenized in PBS and placed in 96-well plate for imaging.

For antitumor assay, 40 tumor-bearing mice were randomly divided into 4 groups (10 per group). Treatments started at the 14^th^ day after tumor inoculation. Treatment was carried out once every 3 days for in total 7 rounds. Drugs were injected via the tail vein. Body wights of mice and tumor volumes were recorded every 3 days. Mice were euthanized once tumor volumes of them above 1200 mm^3^.

### Cell population analysis of tumors

For tumor dissociation, collected tumors were dissociated into a single cell suspension using the Tumor Dissociation Kit, mouse (Miltenyi Biotec, Nordics) in combination with the gentleMACS Octo Dissociator with heaters (Miltenyi Biotec, Nordics) according to the manufacturer’s instructions.

For isolation and counting CAFs via FACS, single cell suspensions of tumors, 200K cells per sample, were suspended in 50 μL of PBS (pH 7.2), 2 mM EDTA, and 0.5% BSA (PEB) buffer. Cy3-labled anti-*α*-SMA antibody was added to cell suspensions and incubated under 4 °C for 10 minutes. After washing via PEB twice, samples were stained with 5 μg/mL propidium iodide (Miltenyi Biotec) immediately before analysis using the MACSQuant^™^ Analyzer (Miltenyi Biotec).

For CD8^+^ T cell analysis in single cell suspensions of tumors, single cell suspensions of tumors were firstly incubated with 12.5 μg/mL mouse IgG (Sigma-Aldrich, USA) PBS for 15 min on ice to block unspecific binding of antibodies. FITC-labeled mouse anti-CD3 antibody and PE-labeled mouse anti-CD8 antibody were diluted in flow buffer consisting of PBS with 10% FBS.

Cell suspensions then were incubated with the antibody mix in 96 v-bottom well plates (Corning, Costar), on ice, in the dark, for 0.5 hour. Following the incubation 100 μL of flow buffer was added to each well, and the plates were centrifuged at 410 g for 6 minutes at 4°C. Supernatants were discarded and cell pellets were re-suspended in 150 μL of flow buffer per well and centrifuged again. Cell viability was assessed by 1 μg/mL propidium iodide prior to flow cytometric analysis. Samples were then analyzed using the MACSQuant™ Analyzer (Miltenyi Biotec).

### Orthotopic model of pancreatic cancer and antitumor assay

We established the orthotopic murine pancreatic cancer model by surgical implantation of 500K Pan02-Luc cells into the head of pancreas of C57BL/6 mice. Treatments, as described above, were started at the 14^th^ day post implant. Bioluminescence imaging was conducted with the IVIS Spectrum system (Caliper Life Sciences, Hopkinton, MA, USA) to monitor the tumor developments. Before the imaging, D-Luciferin (PerkinElmer, USA), dissolved in DPBS, were injected intra-peritoneally to mouse (150 mg D-Luciferin/kg body weight).

### Statistics

All the statistical analysis were carried out in R. When not otherwise stated, results were shown as mean values ± SD. We used two-tailed Student’s t tests to compare data between two groups. We performed one way analysis of variance followed by Turkey posttests to statistically analyze data collected from multiple groups. Significant differences between the groups were indicated by *p < .05, **p < .01, and ***p < .001, respectively.

## ACKNOWLEGMENTS

We thanked Tingting Guo, Ying Yang, Mengling Yao, Ningning Chao and Zhiqiang Liu from Respiratory Health Institute, Frontiers Science Center for Disease-related Molecular Network, West China Hospital, Sichuan University, Chengdu, China and Li Mi from Laboratory of Thyroid and Parathyroid diseases, Frontiers Science Center for Disease-Related Molecular Network, West China Hospital, Sichuan University, Chengdu, China for providing us technique assistance.

## FUNDING

This research is supported by the the National Natural Science Foundation of China (82070625) to T. Wen, the Science and Technology Department of Sichuan Province (2020YJ0237) to Z. Li, the Science and Technology Department of Sichuan Province to W. Wu (2021YFS0230), the 135 project for disciplines of excellence, West China Hospital, Sichuan University (ZYJC18003) to Y. Liu and (2016105 and ZYGD20006) to Z. Zhou, the KWF (Dutch Cancer Society) Young Investigator grant (No. 10140) and a VIDI grant (No. 91719300) from the Netherlands Organisation for Scientific Research (NWO) to Q. Pan.

## AUTHOR CONTRIBUTIONS

Jiaye Liu, MD, PhD (Conceptualization: Lead; Data curation: Lead; Formal analysis: Lead; Methodology: Lead; Project administration: Lead; Writing – review & editing: Lead). Meng Li, PhD (Data curation: Equal; Methodology: Equal; Writing – review & editing: Equal). Kewei Li, MD, PhD (Methodology: Equal; Supervision: Equal; Writing – review & editing: Equal). Yang Wang (Supervision: Lead; Conceptualization: Lead; Formal analysis: Equal; Methodology: Equal; Project administration: Equal; Writing – review & editing: Lead; Funding acquisition: Lead). Shan Li, MD, PhD (Methodology: Equal; Supervision: Equal; Writing – review & editing: Equal). Wenshuang Wu (Supervision: Equal; Writing – review & editing: Equal). Lingyao Du (Conceptualization: Lead; Data curation: Lead; Formal analysis: Lead; Methodology: Lead). Chunyang Mu (Conceptualization: Lead; Data curation: Lead; Formal analysis: Lead; Methodology: Lead). Xiaoyun Zhang (Conceptualization: Lead; Data curation: Lead; Formal analysis: Lead; Methodology: Lead). Chuan Li (Conceptualization: Lead; Data curation: Lead; Formal analysis: Lead; Methodology: Lead). Wei Peng (Conceptualization: Lead; Data curation: Lead; Formal analysis: Lead; Methodology: Lead). Yang Liu (Conceptualization: Lead; Data curation: Lead; Formal analysis: Lead; Methodology: Lead). Dujiang Yang (Data curation: Lead; Formal analysis: Lead; Methodology: Lead; Project administration: Lead). Kaixiang Zhang (Data curation: Lead; Formal analysis: Lead; Methodology: Lead; Project administration: Lead). Qingyang Ning (Data curation: Lead; Formal analysis: Lead; Methodology: Lead; Project administration: Lead). Xiaoying Fu (Data curation: Lead; Formal analysis: Lead; Methodology: Lead; Project administration: Lead). Yu Zeng (Data curation: Lead; Formal analysis: Lead; Methodology: Lead; Project administration: Lead). Yinyun Ni (Data curation: Lead; Formal analysis: Lead; Methodology: Lead; Project administration: Lead). Qiuwei Pan, PhD (Supervision: Lead; Conceptualization: Lead; Formal analysis: Equal; Methodology: Equal; Project administration: Equal; Writing – review & editing: Lead; Funding acquisition: Lead). Zongguang Zhou (Supervision: Lead; Conceptualization: Lead; Formal analysis: Equal; Methodology: Equal; Project administration: Equal; Writing – review & editing: Lead; Funding acquisition: Lead). Yi Liu (Supervision: Lead; Conceptualization: Lead; Formal analysis: Equal; Methodology: Equal; Project administration: Equal; Writing – review & editing: Lead; Funding acquisition: Lead). Yiguo Hu (Supervision: Lead; Conceptualization: Lead; Formal analysis: Equal; Methodology: Equal; Project administration: Equal; Writing – review & editing: Lead; Funding acquisition: Lead). Tianfu Wen, PhD (Conceptualization: Equal; Supervision: Equal; Writing – review & editing: Equal). Zhihui Li, PhD (Conceptualization: Equal; Supervision: Equal; Writing – review & editing: Equal). Yong Liu, PhD (Conceptualization: Equal; Supervision: Equal; Writing – review & editing: Equal).

## CONFLICTS OF INTERESTS

The authors declare that there is no conflict of interests.

## DATA AVAILABILITY

The authors declare that the data that support the findings of this study are available within the paper and its Supplementary Information file. All other information is available from the corresponding authors upon reasonable request.

## Supplementary contents summary

| Items | Contents |
| --- | --- |
| Table S1 | Sequences of DNA barcodes |
| fig. S1 | Zeta potentials |
| fig. S2 | Stability of DMS |
| fig. S3 | AG17724 release from DMS |
| Table S2 | Probed sequences for DNA barcodes hybridization |
| fig. S4 | DNA-antibody conjugates validation |
| Table S3 | Sequence of DNA aptamer targeting CD8<sup>+</sup> T cells |

**Table S1.**
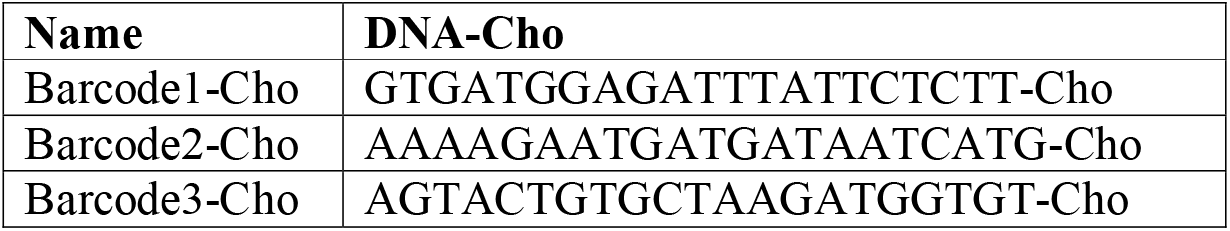
Sequences of DNA barcodes.

**fig. S1.**
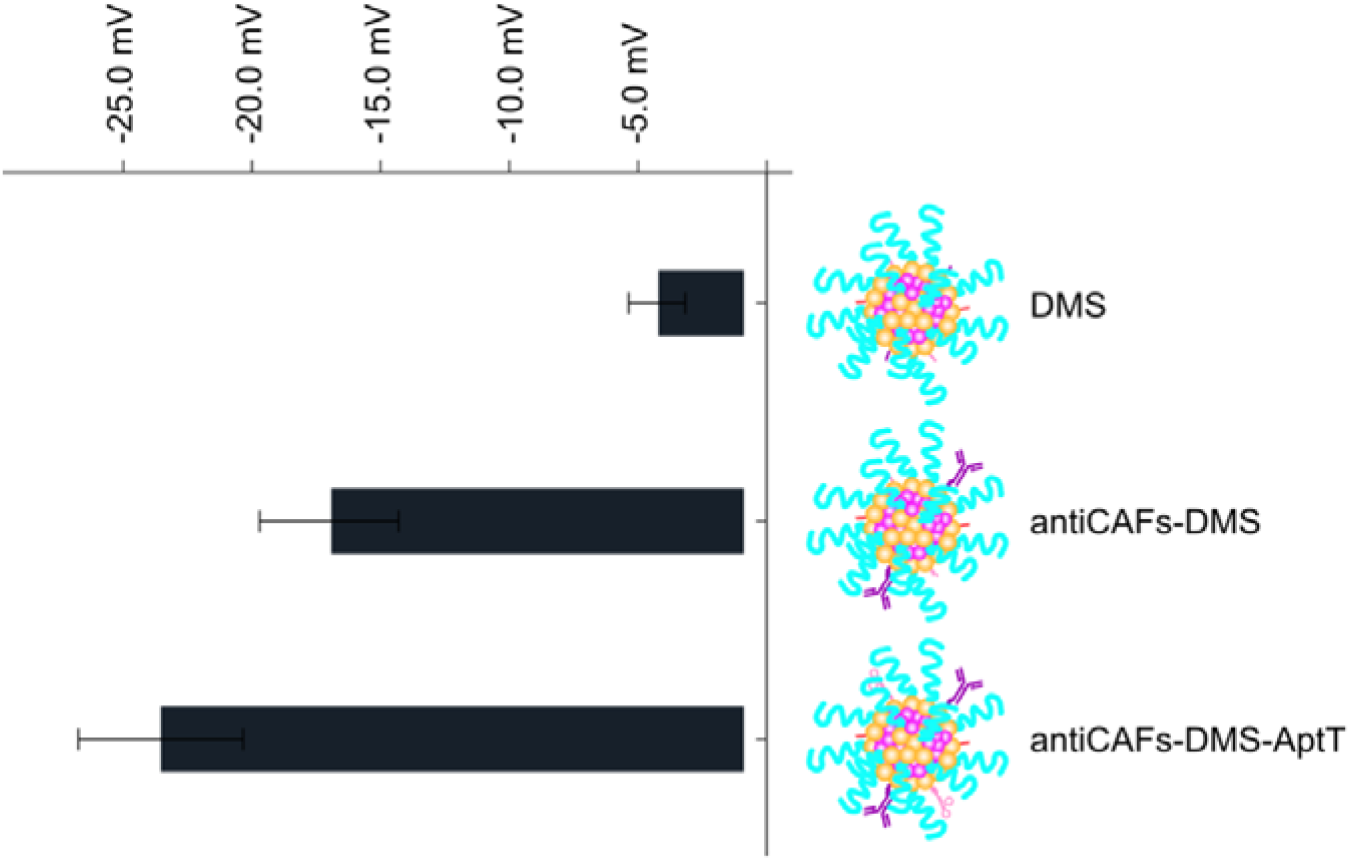
Zeta potentials of DMS, antiCAFs-DMS and antiCAFs-DMS-AptT.

**fig. S2.**
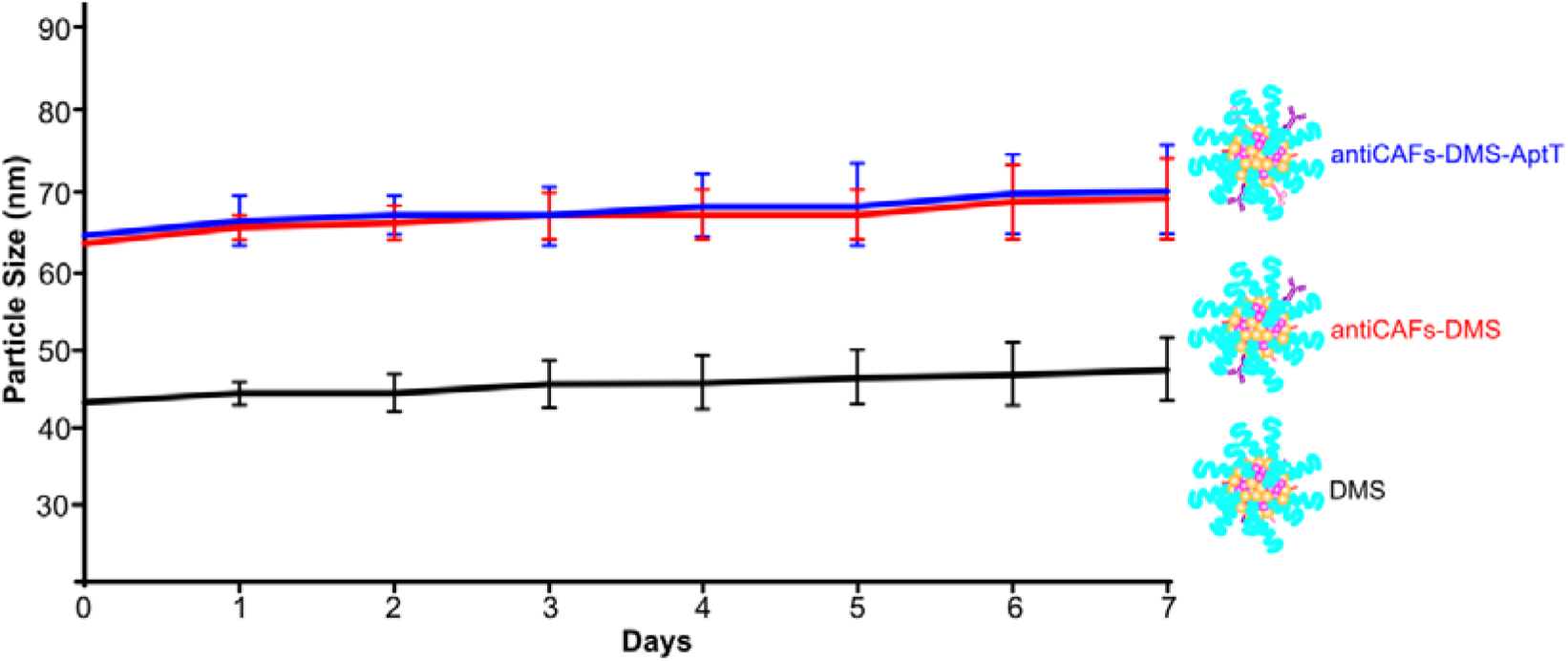
Sizes changes of DMS, antiCAFs-DMS and antiCAFs-DMS-AptT in PBS containing 10% FBS for one week incubation under room temperature.

**fig. S3.**
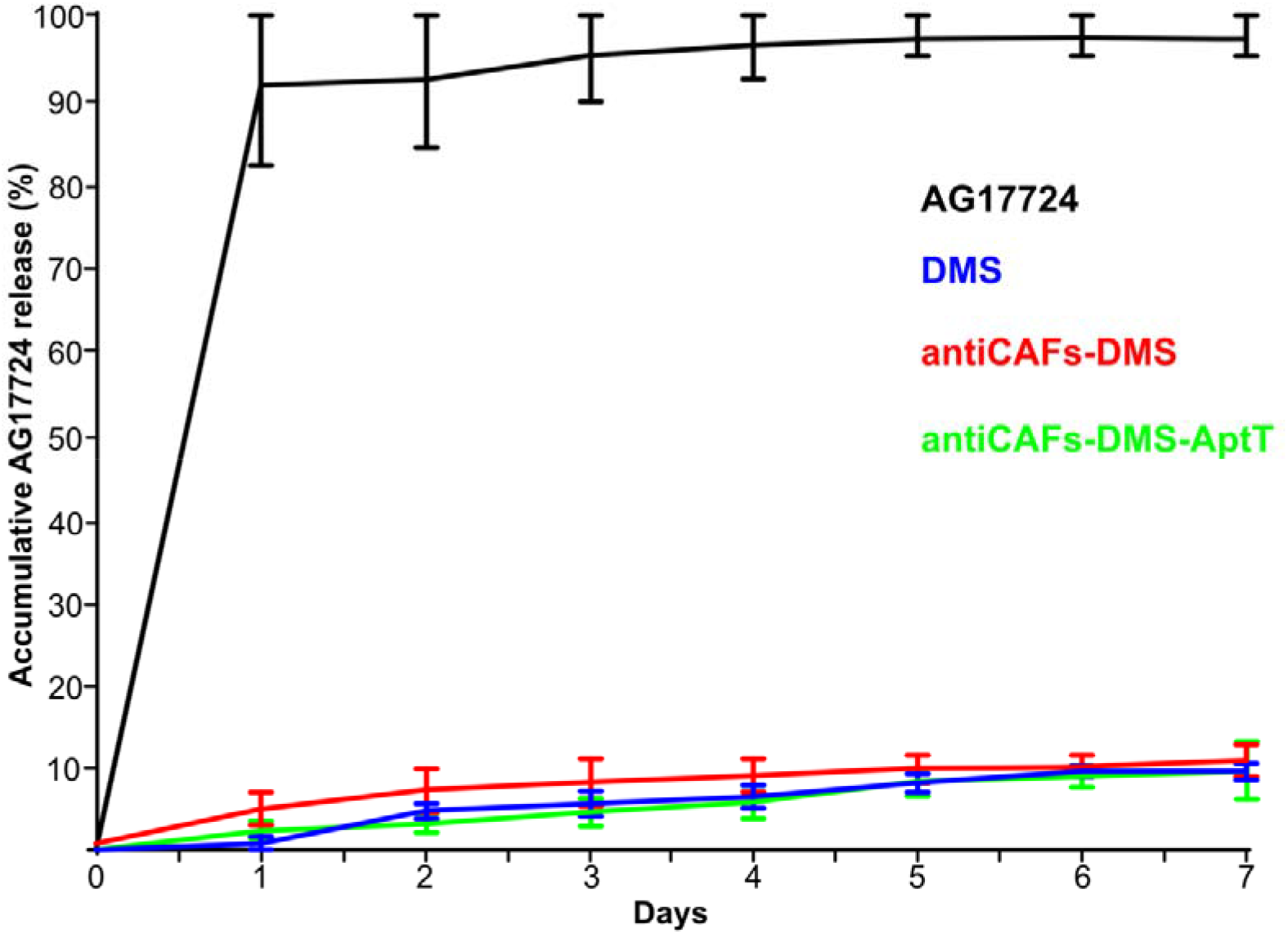
Accumulative release of AG17724 from different preparations in PBS containing 10% FBS for one week under room temperature.

**Table S2.**
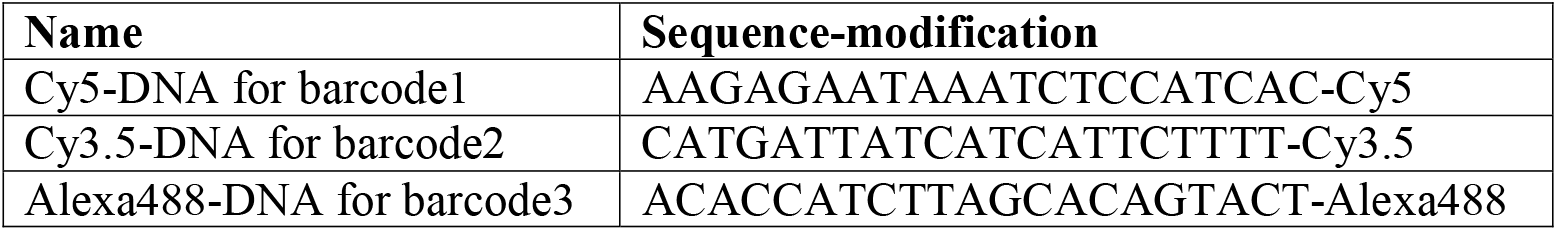
Probed sequences for DNA barcodes hybridization.

**fig. S4.**
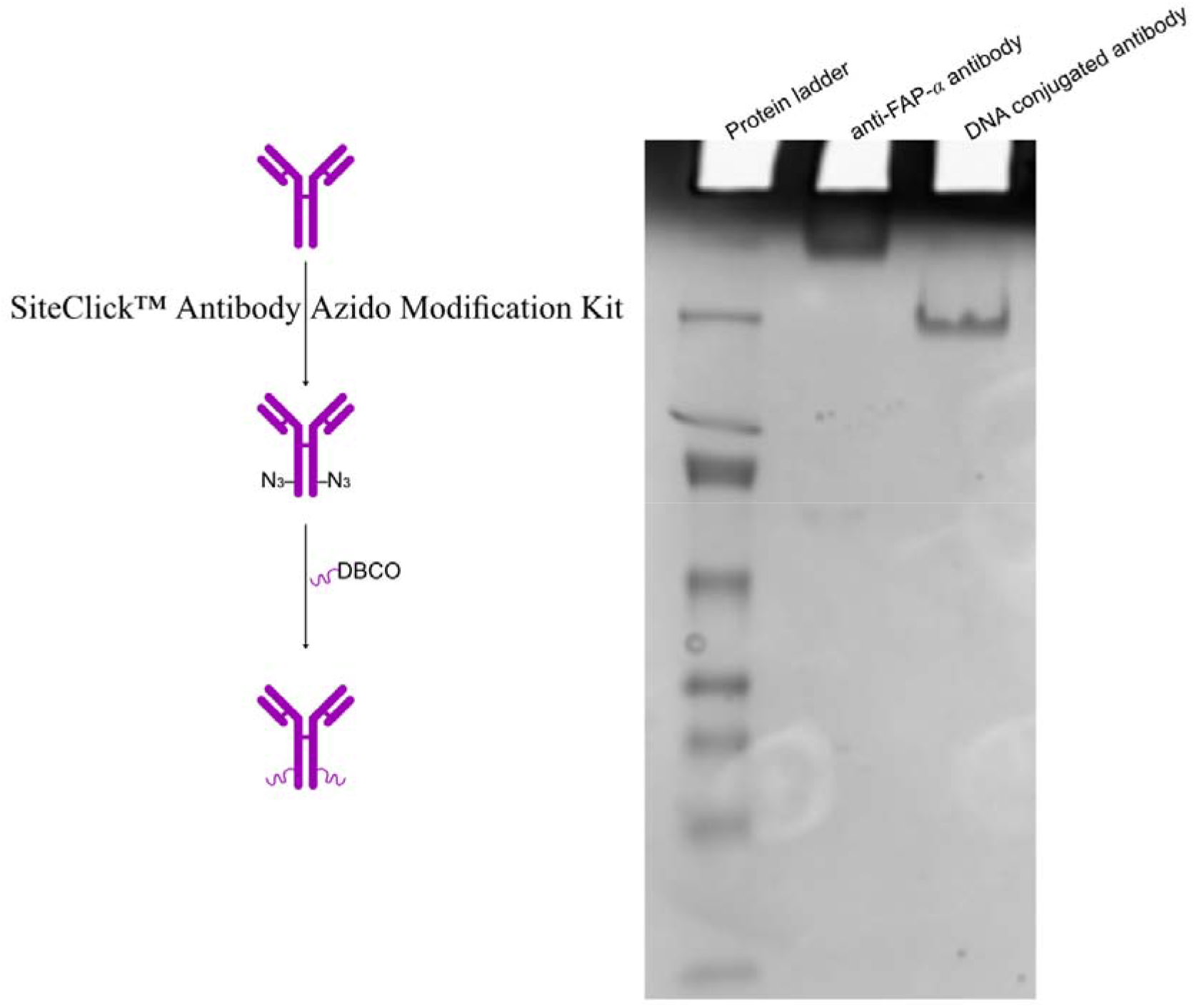
DNA-antibody validation on 10% PAGE gel. The gel was stained with Coomassine.

**Table S3.**
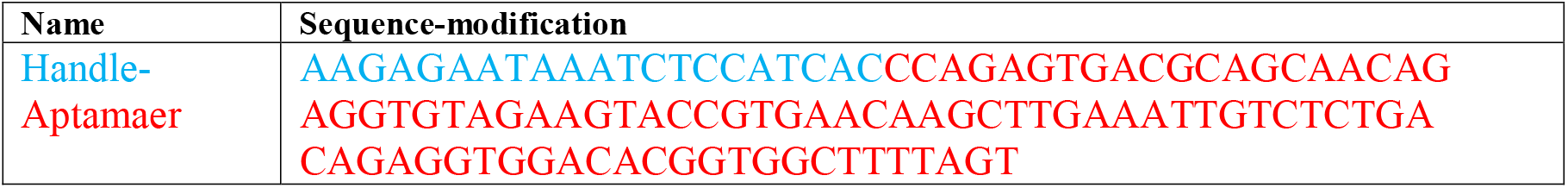
Sequence of DNA aptamer targeting CD8^+^ T cells.

## Notes

### Competing Interest Statement

The authors have declared no competing interest.

